# Rewiring the carbon cycle: a theoretical framework for animal-driven ecosystem carbon sequestration

**DOI:** 10.1101/2023.07.14.549071

**Authors:** Matteo Rizzuto, Shawn J. Leroux, Oswald J. Schmitz

## Abstract

Most carbon cycle models do not consider animal-mediated effects, focusing instead on carbon exchanges among plants, microbes, and the atmosphere. Yet, a growing body of empirical evidence from diverse ecosystems points to pervasive animal effects on ecosystem carbon cycling and shows that ignoring them could lead to misrepresentation of an ecosystem’s carbon cycle. We develop a new theoretical framework to account for animal effects on ecosystem carbon cycling. We combine a classic ecosystem compartment modeling approach with a classic carbon model to account for carbon flux and storage among plant, animal, and soil microbial trophic compartments. We show, by way of numerical analyses of steady state conditions, that herbivore presence alters the dominant pathways of control over carbon storage and capture. This altered control arises via direct, consumptive effects and especially via indirect, non-consumptive pathways by instigating faster nutrient recycling. This leads to a quantitative change in the ecosystem’s carbon balance, increasing the amount of carbon captured and stored in the ecosystem by 2–3 fold. The modeling shows that animals could play a larger role in ecosystem carbon cycle than previously thought. Our framework provides further guidance for empirical research aimed at quantifying animal-mediated control of carbon cycling and to inform the development of nature-based climate change solutions that leverage animal influence on the carbon cycle to help mitigate climate change.

## 1 Introduction

The carbon cycle involves a series of biogeochemical processes that move carbon through plant, animal, and microbial trophic compartments within ecosystems, and between ecosystems and the atmosphere. The series of processes fundamentally includes photosynthetic capture and assimilation of atmospheric carbon into plant biomass, consumptive transfer and assimilation of carbon into animal (herbivore and carnivore) biomass, assimilation of detrital plant and animal biomass carbon into microbial biomass, and autotrophic (plant) and heterotrophic (microbial and animal) respiratory release of carbon to the atmosphere (Keenan and Williams 2018; Middelburg 2019). Thus, all plants, animals, and microbes contribute toward regulating biomass carbon storage in ecosystems (Steinberg and Landry 2017; Middelburg 2019; Schmitz and Leroux 2020). Yet the predominance of carbon cycle models only account for carbon movement and storage among plant and microbial trophic compartments and the atmosphere (Piao et al. 2013; Zaehle et al. 2014).

Such model formalism is based on two tacit assumptions: (i) ecosystem carbon capture and storage is controlled only by nutrient and water limitations (i.e., ecosystem processes are bottom-up controlled); and (ii) animal contributions are negligible, or at least are so miniscule that they can be safely subsumed by effects of larger biomass compartments, e.g., animal respiration is overwhelmed by microbial respiration (Schmitz and Leroux 2020; Rastetter et al. 2022). However, such assumptions can mischaracterise how ecosystem carbon cycling is controlled, in turn, potentially leading to large under- or over-estimates of the amount of carbon storage in ecosystems (Schmitz and Leroux 2020; Schmitz, Sylvén, et al. 2023).

Here we illustrate how including animal feedback controls (i.e., top-down control of ecosystem processes) in a carbon cycle model can fundamentally change the importance of different pathways and rates of carbon movement (fluxes) among trophic compartments and the atmosphere, relative to a model that excludes animal controls. The inclusion of animals in carbon cycle models is rare, but growing (Huntley, Lopez, and Karl 1991; Leroux, Hawlena, and Schmitz 2012; Metcalfe et al. 2014; Leroux and Schmitz 2015; Dangal et al. 2017; Rastetter et al. 2022). However, these earlier models are typically designed to characterize specific empirical systems, thereby limiting generalization. We instead designed our models to provide a scaffolding for more generalizable analyses (*sensu* Ives, Cardinale, and Snyder 2005; Leroux and Schmitz 2015). As such, our process-based modeling does not contain the details needed to depict any specific, real-world system, because such empirical details, while growing, are still lacking for most systems (Schmitz and Leroux 2020; Pringle et al. 2023). The model does, however, contain the salient principles of animal control over biogeochemical processes (aka, zoogeochemistry; Schmitz, Wilmers, et al. 2018) that apply broadly across ecosystems. Furthermore, the structure of the model does enable the substitution of functions for parameters to mechanistically characterize processes and interactions that apply to a given real-world system. By doing this, we aim to inspire new quantitative empirical measurements of animal effects in all kinds of ecosystem types. Considering animal effects on ecosystem processes will be important because, as we show that animal effects could be disproportionately larger than expected based merely on their biomass representation within ecosystems, thereby counterbalancing bottom-up control. This arises because animals cause direct and indirect feedback effects that alter the abundance, elemental content and biogeochemical functioning of the larger, plant and microbial biomass pools.

## 2 Model Structure

Our model combines a classic ecosystem compartment model (DeAngelis 1992; Holt 1997; Loreau 2010) with a classic carbon budget model (Chapin, Woodwell, et al. 2006; Ballantyne et al. 2021). The modeling uses principles of elemental flux and storage among different trophic levels in an ecosystem, based on known processes for their action (Leroux and Loreau 2010). The ecosystem trophic compartment structure—which includes soil elemental pools, plants, and animals (Figure 1a, c)—is typically used when examining organismal effects on ecosystem functioning (Hall et al. 2007; Leroux and Loreau 2010; Loreau 2010; Bassar et al. 2012; Leroux, Hawlena, and Schmitz 2012). We use a stoichiometric approach that focuses on fluxes and pool sizes of nitrogen (N) and carbon (C) to account for rate limitation of carbon cycling by an essential nutrient. We focus on C and N because they (i) share a strong stoichiometric relationship captured by the C:N ratio which describes the stoichiometry of an individual or trophic/functional group and allows for tracking both C and N (Sterner and Elser 200; Schmitz and Leroux 2020), (ii) are fundamental actors in existing carbon models (Zaehle et al. 2014), and (iii) are routinely measured in organisms as part of empirical studies (Meunier et al. 2017; Welti et al. 2017). Nonetheless, this model formalism does not preclude the inclusion or substitution of other nutrients (e.g., phosphorus) that may limit carbon cycling rate.

We first describe an ecosystem model tracking both C and N but without herbivores, and then build on this scaffold to introduce herbivores in the model. These two models capture the essential features of elemental cycling in general (DeAngelis 1992; Loreau 2010) and carbon cycling specifically (Schmitz and Leroux 2020), including elemental uptake by plants from the abiotic environment (i.e., C uptake from the atmosphere and N uptake from the soil elemental pool) and elemental transfer to and loss from all compartments through trophic interactions, respiration, excretion, egestion, and leaching out of the ecosystem. As such, the models depict open systems, i.e., elements are not solely recycled within the confines of the ecosystem. Nevertheless, they are formulated to obey fundamental mass balance requirements (Loreau 2010) such that, at equilibrium, elemental inputs to the ecosystem equal elemental losses from the ecosystem plus storage. After describing our ecosystem compartment model without and with herbivores, we then outline how we investigate the impacts of herbivores on ecosystem structure, function and net ecosystem carbon balance.

### 2.1 Model without Herbivores

Our base model without herbivores (Figure 1a and Equations (1a) to (1d)) comprises two trophic compartments: the soil (𝑆_𝑗_), containing the inorganic nutrient pools, and the primary producers (i.e., plants; 𝑃_𝑗_), where the subscript 𝑗 ∈ [𝐶, 𝑁]. Following similar ecosystem formulations (Leroux and Loreau 2008; Loreau 2010) we do not explicitly model detritus, decomposition or mineralization/immobilization processes which animals also may influence (Kristensen et al. 2022; Naidu, Roy, and Bagchi 2022; Losada et al. 2023); rather, our scaffold implicitly accounts for these ecosystem components and processes as part of the soil compartment (𝑆_𝑗_) as a starting approximation. Functions to mechanistically characterize these processes can be incorporated using data available for a given real-world system. The current model formalism assumes that recycled organic materials are converted to inorganic form. Inorganic N enters the system at rate 𝐼 and leaches out from the soil compartment at rate 𝑘. 𝛷_𝑃_ represents plant consumption of N, while 𝜃_𝑃_ captures N losses from the plant compartment. We assume that all N lost from the plants is recycled back into the soil (Gravel et al. 2010). Parameter 𝛼 captures the stoichiometric relationship between C and N in plants (Sterner and Elser 2002), thus binding the two elements in the model. Parameter 𝛿 represents the rate of C loss via plant respiration. Thus, the terms 𝛼𝛷_𝑃_ and 𝛼𝛿𝜃_𝑃_ on the C side of the model capture plant photosynthesis and respiration, respectively, and are the main pathways for C to enter and leave the system (Figure 1a and Table 1). The term (1 − 𝛿)𝛼𝜃_𝑃_ captures the fraction of C from the plant compartment that is recycled back into the soil. From the soil, C leaves the system at respiratory rate 𝛾_𝑆_. The following set of ordinary differential equations (ODE) describes the model ecosystem without herbivores that embodies these processes:

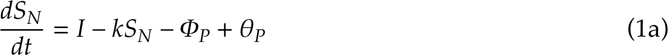

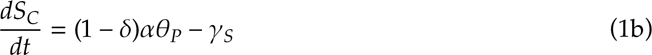

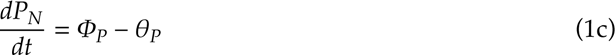

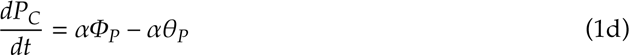

**Figure 1:**
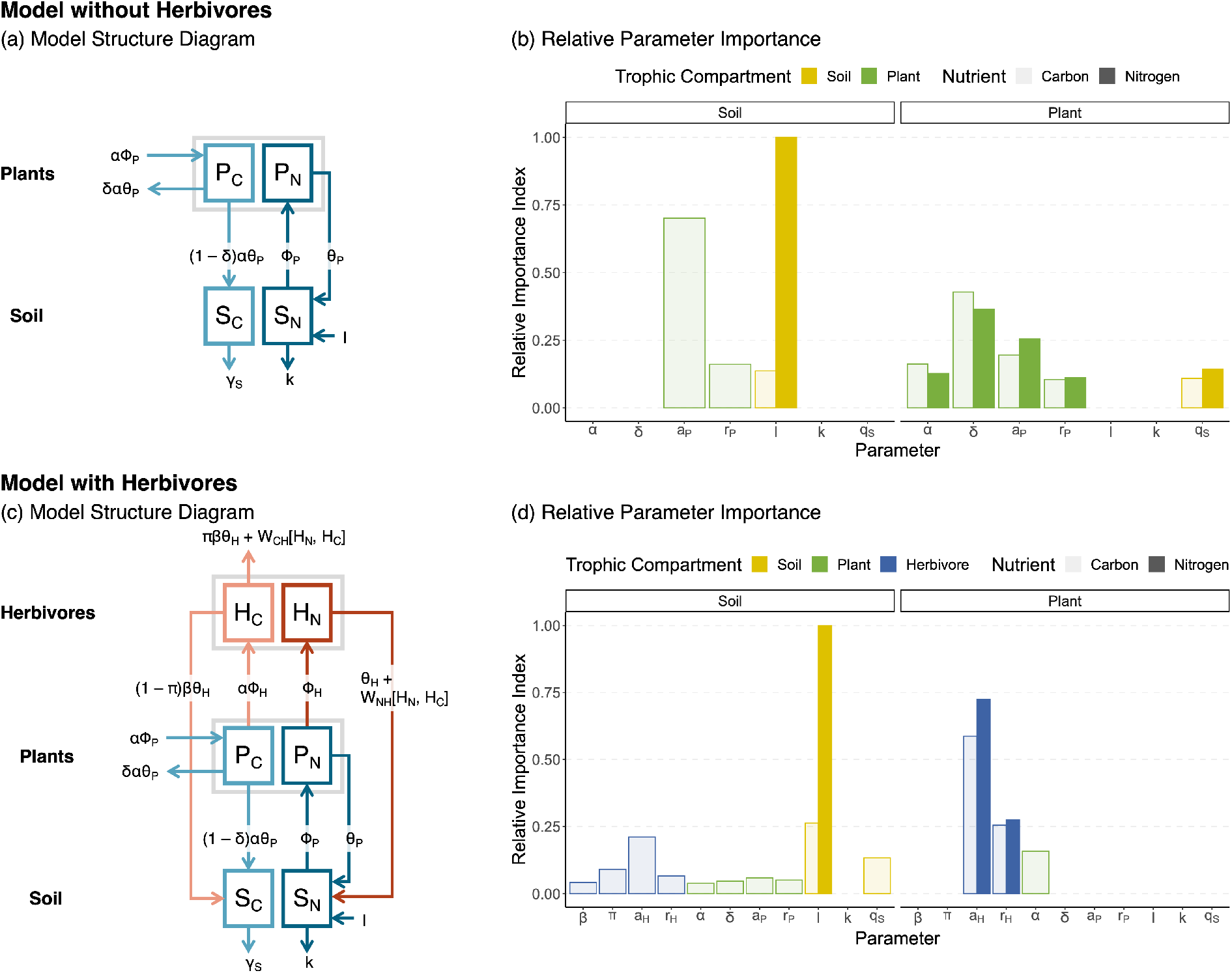
Adding herbivores to a simple plant-soil model tracking both carbon and nitrogen (panels (a) and (c)) results in a rewiring of the trophic food web (compares panels (b) and (d)). In particular, the parameters that capture the herbivores’ growth (𝑎_𝐻_) and recycling (𝑟_𝐻_) have an out-sized influence on the dynamics of the plant compartment for both nutrients tracked by the model. Herbivores’ respiration (𝜋) appears to influence carbon dynamics in the soil to a larger extent than plant respiration. As well, herbivores’ presence cancels the effect of plant C:N (𝛼) on plant N stocks, but this parameters maintains influence on plant C stocks. In panels (a) and (c), boxes and arrows in shades of blue identify the the plant-soil system, while herbivore addition is shown in shades of red. In panels (b) and (d), we group parameters on the y-axis by the trophic compartment they belong (Equations (1a) to (2f)), as shown by the different colors of the bars. Different shades of the same color represent the relative importance of a given parameter for carbon (light) and nitrogen (solid).

**Table 1:** Model functions descriptions and parameter definitions. See Table 2 for units of measurement, as well as further parameter definitions and value ranges. Abbreviations: N, nitrogen; C, carbon; S, soil; P, plants; H, herbivores. Appendix B reports the derivation of the 𝑊_𝐶𝐻_[𝐻_𝑁_, 𝐻_𝐶_] term.

| Functions | Description | Formula | Parameters & Definitions |
| --- | --- | --- | --- |
| $\Phi_P$ | N flux from soil to plant compartment | $a_P P_N S_N S_C$ | $a_P$, N mineralization rate |
| $\Phi_H$ | N flux from plant to herbivore compartment | $a_H P_N H_N$ | $a_H$, herbivore nutrition rate |
| $\theta_i$ | recycling flux from trophic compartment $i$ to the soil | $r_i i_N$ | $r_i$, N-recycling rate of trophic level $i$, where $i \in [P, H]$ |
| $\gamma_S$ | inorganic loss rate of C from soil | $q_S S_N S_C$ | $q_S$, C loss rate from soil compartment |
| $W_{CH}[H_N, H_C]$ | herbivore differential assimilation rate of C | $(\alpha - \beta) a_H P_N H_N$ | $\alpha$, plant C:N ratio; $\beta$, herbivore C:N ratio |

Table 1 provides definitions and formulae for functions 𝛷_𝑃_, 𝜃_𝑃_, and 𝛾_𝑆_, and Table 2 provides definitions of all parameters.

**Table 2:** Variables and parameters of the model shown in Figure 1a, c. Abbreviations: N, nitrogen; C, carbon.

| Variables and Parameters | Symbol | Units | Range | Notes |
| --- | --- | --- | --- | --- |
| Soil pool of nutrient $j$ | $S_j$ | $g$ | $> 0$ | where $j \in [N, C]$ |
| Plant pool of nutrient $j$ | $P_j$ | $g$ | $> 0$ | where $j \in [N, C]$ |
| Herbivore pool of nutrient $j$ | $H_j$ | $g$ | $> 0$ | where $j \in [N, C]$ |
| Soil inorganic input rate of N | $I$ | $g \times t^{-1}$ | $> 0$ | |
| Soil inorganic loss rate of N | $k$ | $g \times t^{-1}$ | $> 0$ | |
| Soil inorganic loss rate of C | $\gamma_S$ | $g \times t^{-1}$ | $> 0$ | |
| C:N ratio of plants | $\alpha$ | unit-less | $> 0$ | Under N-limitation, $\alpha > \beta$ |
| C:N ratio of herbivores | $\beta$ | unit-less | $> 0$ | Under N-limitation, $\alpha > \beta$ |
| Proportion of plant C that is respired | $\delta$ | unit-less | $[0, 1]$ | $1 - \delta$ is the proportion of C recycled back into the soil |
| Proportion of herbivores C that is respired | $\pi$ | unit-less | $[0, 1]$ | $1 - \pi$ is the proportion of C recycled back into the soil |
| N mineralization rate | $a_P$ | $g \times t^{-1}$ | $> 0$ | see Table 1 |
| Herbivore nutrition rate | $a_H$ | $g \times t^{-1}$ | $> 0$ | see Table 1 |
| N-specific recycling rate of trophic level $i$ | $r_i$ | $g \times t^{-1}$ | $> 0$ | where $i \in [P, H]$; see Table 1 |
| C loss rate from the soil | $q_S$ | $g \times t^{-1}$ | $> 0$ | see Table 1 |

### 2.2 Model with Herbivores

We build on the foundational model (equations (1a) to (1d)) to add a herbivore consumer trophic compartment to both sets of equations specifying C and N dynamics. In the following system of ODE, 𝐻_𝑗_ is the herbivore compartment (𝑗 ∈ [𝐶, 𝑁]; Figure 1c and equations (2a) to (2f)). The term 𝛷_𝐻_ captures the herbivores’ consumptive N uptake from the plant compartment, while 𝜃_𝐻_ represents herbivores’ N losses via excretion and carcass deposition. 𝛽 represents the stoichiometric C:N ratio of herbivores, and 𝜋 captures their rate of gaseous C release (which can flexibly include respiratory release as CO_2_ and enteric release as CH_4_). Hence, the terms 𝜋𝛽𝜃_𝐻_ and (1 − 𝜋)𝛽𝜃_𝐻_ capture the herbivores’ release of C to the atmosphere and the recycled fraction of C released as egesta from herbivores to the soil compartment, respectively. Finally, the term 𝑊_𝐶𝐻_[𝐻_𝑁_, 𝐻_𝐶_] represents the fraction of assimilated C that herbivore release over time as a function of their body content of C and N (Daufresne and Loreau 2001a,b).

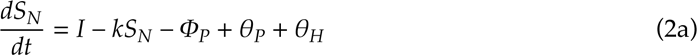

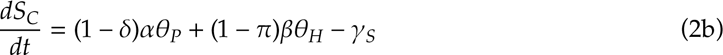

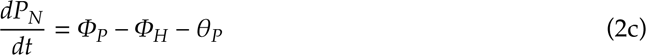

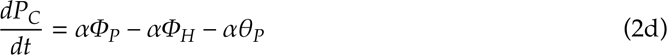

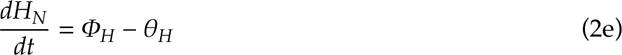

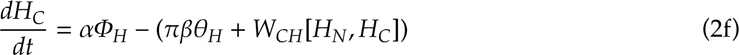

Terms and parameters pertaining to the Soil (𝑆_𝑗_) and Plants (𝑃_𝑗_) compartments remain the same as in the model without herbivores (Equations (1a) to (1d)). As with the model without herbivores, we assume that all N lost from plants and herbivores is recycled back into the soil compartment (Gravel et al. 2010). However, unlike earlier models, our model does not include a parameter that quantifies trophic transfer efficiency directly. Rather, transfer efficiency becomes an emergent consequence of physiological first principles accounting of C gains due to consumption (parameter 𝑎_𝐻_; Table 2), and C losses from gaseous release (parameter 𝜋; Table 2) and egesta release (the term (1 − 𝜋)𝜃_𝐻_ in equation (2b)) from the herbivore compartment. See Table 1 for a description of each function in the model, and Table 2 for a description of all model parameters. Appendix A reports equilibria derivation, as well as feasibility conditions, and Appendix B details the derivation of the 𝑊_𝐶𝐻_[𝐻_𝑁_, 𝐻_𝐶_] term.

## 3 Numerical Analyses

We solved equations (1a) to (1d) and equations (2a) to (2f) for equilibria (mass balance) in Mathematica (v.13.2; Wolfram Research 2022). Each model has one equilibrium where all state variables can be positive, and thus biologically feasible (see Appendix A). We use these equilibria for the model *without* and *with* herbivores to perform further numerical analyses in R (v. 4.3.1; R Core Team 2022) aimed at assessing how different magnitudes of model parameter values influence C stocks in biomass pools, net primary productivity (net carbon capture by plants) and net ecosystem carbon balance (the net amount of carbon stored within the ecosystem). We further characterized the sensitivity of the models to changes in model structure and parameters. Following Leroux and Schmitz (2015), we used a latin hypercube sampling design to generate 10 000 sets of random parameter values for all parameters in the models, using function randomLHS from the lhs R package (Carnell 2022). We scaled the random parameter values assigned to parameters 𝛼 and 𝛽, representing the C:N ratio for plants and herbivores, respectively, to fit in the empirical C:N ranges reported in Buchkowski, Leroux, and Schmitz (2019). As well, for parameters 𝛿 and 𝜋, representing the respiration rates for plants and herbivores respectively, we rescaled the randomly drawn parameter values to be ∈ [0, 1]. We applied the parameter sets to both models, *without* and *with* herbivores. Of the 10 000 iterations, we retained only those model solutions that satisfied the equilibria’s feasibility conditions (i.e., 𝑆_𝑗_^∗^, 𝑃_𝑗_^∗^, and 𝐻_𝑗_^∗^ > 0, where 𝑗 ∈ [𝐶, 𝑁]; reported in Appendix A).

### 3.1 Ecosystem structure analyses

We performed a global sensitivity analysis (Harper, Stella, and Fremier 2011; Bellmore et al. 2014) to evaluate how each parameter (Table 2) influences the performance of either model. A global sensitivity analysis (GSA) varies the values of all parameters simultaneously, thereby accounting for parameter interactions when quantifying each parameter’s influence on the models’ results (Harper, Stella, and Fremier 2011). Using a GSA allows us to (i) investigate changes in the relative influence of parameters in the two models and (ii) quantify each parameter’s effect in shaping the three ecosystem functions we use to investigate herbivore effects on ecosystem functioning and C sequestration (see below). We use function randomForest in R to fit a random forest algorithm to the feasible parameter sets (see above) used in our numerical analyses. We use as response variables the values of each state variable at equilibrium and for each ecosystem function of interest when fitting the random forest algorithm. Function randomForest computes the residual sum of squared errors (SSE) for each parameter; from which we calculate a normalized index of relative importance by dividing each parameter’s SSE by the total SSE. We repeated this process for both the model *without* and *with* herbivores. Finally, we use paired bar plots to visually assess how each parameter influences the model results in each scenario (*without* vs *with* herbivores; see the Supplementary Code document available in the online repository; Rizzuto, Leroux, and Schmitz 2023).

### 3.2 Ecosystem function analyses

We investigate the effects of herbivore presence and absence on carbon cycling by focusing on three ecosystem functions: biomass and stock accumulation, primary productivity, and net ecosystem carbon balance (NECB; Chapin, Woodwell, et al. 2006; Schmitz, Sylvén, et al. 2023). We use the equilibrium values 𝑆∗_𝑗_ and 𝑃_𝑗_^∗^ as estimates of equilibrium soil stock and plant biomass accumulation (g), where 𝑗 ∈ [𝐶, 𝑁]. Further, following Loreau (2010), we estimate ecosystem primary productivity at equilibrium, by substituting the values of 𝑆_𝑗_^∗^, 𝑃_𝑗_^∗^, and 𝐻_𝑗_^∗^ in the expressions for 𝛷_𝑃_ (Table 1). Finally, we estimate NECB by explicitly accounting for the productivity of each of plant and herbivore trophic compartments (see below for details). We compared the values of these three ecosystem functions at equilibrium between the herbivore and non-herbivore model to investigate the effects of adding herbivore to the model ecosystem. As well, we use partial derivatives of the equilibria with respect to the herbivores’ attack rate on plants (𝑎_𝐻_) and nutrient recycling rate (𝑟_𝐻_) to evaluate how trophic processes to capture and store atmospheric C are mediated by herbivore consumers.

### 3.3 Net Ecosystem Carbon Balance derivation

Net Ecosystem Carbon Balance (NECB) measures ecosystem carbon storage in terms of the net difference between an ecosystem’s anabolic and catabolic processes, i.e., the balance between net rate of carbon accumulation in ecosystems due to carbon fixation by plants (primary productivity), heterotrophic production, heterotrophic respiratory CO_2_ emissions, as well as additional losses including CH_4_ emissions directly from animals and soils and sediments of ecosystems (Chapin, Matson, and Vitousek 2002; Chapin, Woodwell, et al. 2006). Traditionally, net ecosystem carbon balance is estimated as

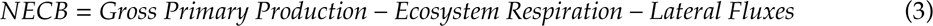

in which Gross Primary Production is essentially gross carbon uptake by plants, Ecosystem Respiration comprises respiration by all trophic compartments (e.g., soil, plants, herbivores; Chapin, Woodwell, et al. 2006)), and Lateral Fluxes comprise losses of C through mechanisms other than respiration (e.g., leaching to groundwater, methane emissions Chapin, Matson, and Vitousek 2002).

However, by only debiting the respiration of heterotrophs from autotroph (primary) production, equation (3) does not account for the direct and indirect interactions among an ecosystem’s trophic compartments—net assimilation of C in animal biomass (secondary productivity that varies with animal stoichiometry) and N and C release from animals (recycling feedbacks that also vary with animal stoichiometry) that promote Gross Primary Production—thereby missing important contributions to an ecosystem’s C budget by these actors via the processes they mediate.

Hence, we expand on equation (3) to integrate the effects of heterotrophs in the accounting of NECB. Broadly, we define NECB as the sum of Net Primary Production (NPP) and Net Heterotrophic Production (NHP):

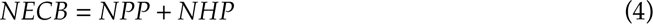

These two components capture the combined anabolic and catabolic processes happening in an ecosystem, across all trophic compartments. Net Primary Production is the balance of the photosynthetic and respiratory processes that happen in the autotroph compartment of an ecosystem. If we imagine a terrestrial ecosystem, where primary producers are generally plants, we have that:

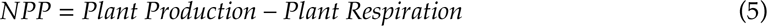

Conversely, Net Heterotrophic Production is the algebraic sum of all biomass-producing and respiratory processes taking place in the heterotrophic compartments of the ecosystem. These include (i) any trophic level above the autotrophs—e.g., herbivores, predators—but also (ii) any trophic level involved in the decomposing pathways that recycle nutrients from waste and dead biomass and make them available to autotrophs once again—the so-called “brown food web”. In our case (Figure 1c), (i) is the herbivores and (ii) is the soil. So, using the same terrestrial ecosystem example as equation (5), we obtain:

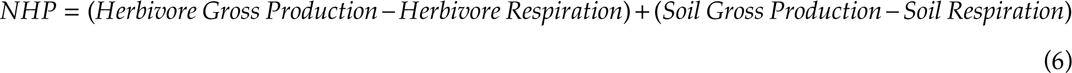

where Gross Production is measured as C uptake rate by the herbivore and soil trophic compartments. Note that equation (6) can accommodate more complex model formulations, for instance, models with trophic compartment for a predator, detritus/decomposer, or both. Equation (4) is conceptually comparable to equation (3), but allows for debiting heterotroph respiration from heterotroph production. Accounting for both components of heterotrophic carbon effects allows us to explicitly measure the relative impact of different kinds of heterotrophs on NECB. When substituting the equilibrium values of the relevant functions and state variables into equations (5) and (6), the model without herbivores specifies𝑁𝐸𝐶𝐵_¬𝐻_ as:

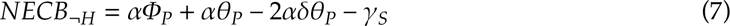

and the model with herbivores specifies NECB as:

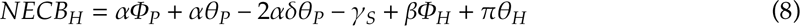

Appendix C presents the full derivation of equations (7) and (8).

## 4 Results

Our results arise from conducting numerical estimates of ecosystem functions that produce large sample sizes of output due to an ensemble of parameter selections. We therefore refrain from comparing treatments (herbivore absent vs. present) using frequentist statistical tests on the data because such data produce artificially low *p*-values and risk committing a Type II statistical error (White et al. 2014). We follow recommendations to report effect sizes of differences and use graphical means to explore and compare the results arising from the herbivore and non-herbivore versions of our model (White et al. 2014).

### 4.1 Herbivore presence rewires the ecosystem’s trophic pathways

Introducing herbivores into the ecosystem model (Figure 1a vs 1c) leads to an extensive rewiring of trophic pathways, consistent with a shift from pervasive bottom-up to top-down control (Figure 1b, d). This trophic rewiring influences the dynamics of both elements, C and N, tracked by the model (Figure 1b, d). In the model without herbivores (Figure 1b), results of the global sensitivity analysis align with insights from conventional carbon cycle models in that primary producers dominate carbon dynamics of both soil and plant trophic compartments through consumption (parameter 𝑎_𝑃_), respiration (𝑟_𝑃_), and respiration (𝛿). Moreover, inorganic inputs to the soil compartment (𝐼) have a disproportionately large influence on N dynamics compared to all other parameters, underscoring N-limitation of ecosystem dynamics. The model with herbivores (Figure 1d) results in an important rearrangement of which parameters drive ecosystem stocks and fluxes of C and N. The herbivores’ attack rate (𝑎_𝐻_) becomes the leading driver of C accumulation in the plant compartment, and the second most important determinant of C in the soil compartment after the input of inorganic N into the system (Figure 1d). As well, C loss from the soil compartment (𝑞_𝑆_) appears to matter less for plant and more for soil C dynamics when herbivores are present in the system than when they are absent. For N, soil compartment dynamics are still strongly controlled by external inputs (𝐼). In plants, however, N dynamics in the presence of herbivores become dominated by parameters regulating the herbivores’ functional response—i.e., the herbivores’ attack (𝑎_𝐻_) and recycling (𝑟_𝐻_) rates (see Table 2 for definitions). This top-down control of trophic pathways in the model with herbivores results in equilibrium N accumulation resembling a classic trophic cascade (Figure D.2). At the same time, the system is sequestering larger quantities of C from the atmosphere and storing them in biomass—either the plants’ or the herbivores’—compared to a system devoid of herbivores (Figures D.1 and D.2).

### 4.2 Herbivore-increased biomass turnover and recycling boost primary productivity

Increased atmospheric C sequestration by the ecosystem with herbivores present appears likely also due to the two-fold increase in primary productivity in this system compared to the ecosystem model without herbivores (Figure 2a). Again, this increase is mediated by the herbivores’ functional response parameters—attack (𝑎_𝐻_), recycling (𝑟_𝐻_), and respiration (𝜋) rates—which dampen the effect of the plants’ own parameters involved in consumption and growth (Figure 2b). Notably, plant and herbivores C:N stoichiometry influence primary productivity when herbivores are present, whereas they do not when herbivores are absent (solid-color bars for parameters 𝛼 and 𝛽 in Figure 2b). Partial derivatives of primary productivity with respect to the herbivores’ attack and recycling rates (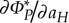 and 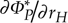, respectively) provide some insights in how their influence plays out. The herbivores’ attack rate has an overall dampening effect on primary productivity (Figure 2c), causing primary productivity to fall for the vast majority of parameter values explored by our model. Some values of 𝑎_𝐻_, however, have the opposite effect and lead to an increase in primary production (shown in red and highlighted in the inset in Figure 2c). Conversely, herbivores recycling rate 𝑟_𝐻_ substantially increases primary productivity, leading to an overall increase in the plant’s productivity (Figure 2d). Again, some of the values of this parameter explored in our model have the opposite, negative effect (shown in red and highlighted in the inset in Figure 2d).

**Figure 2:**
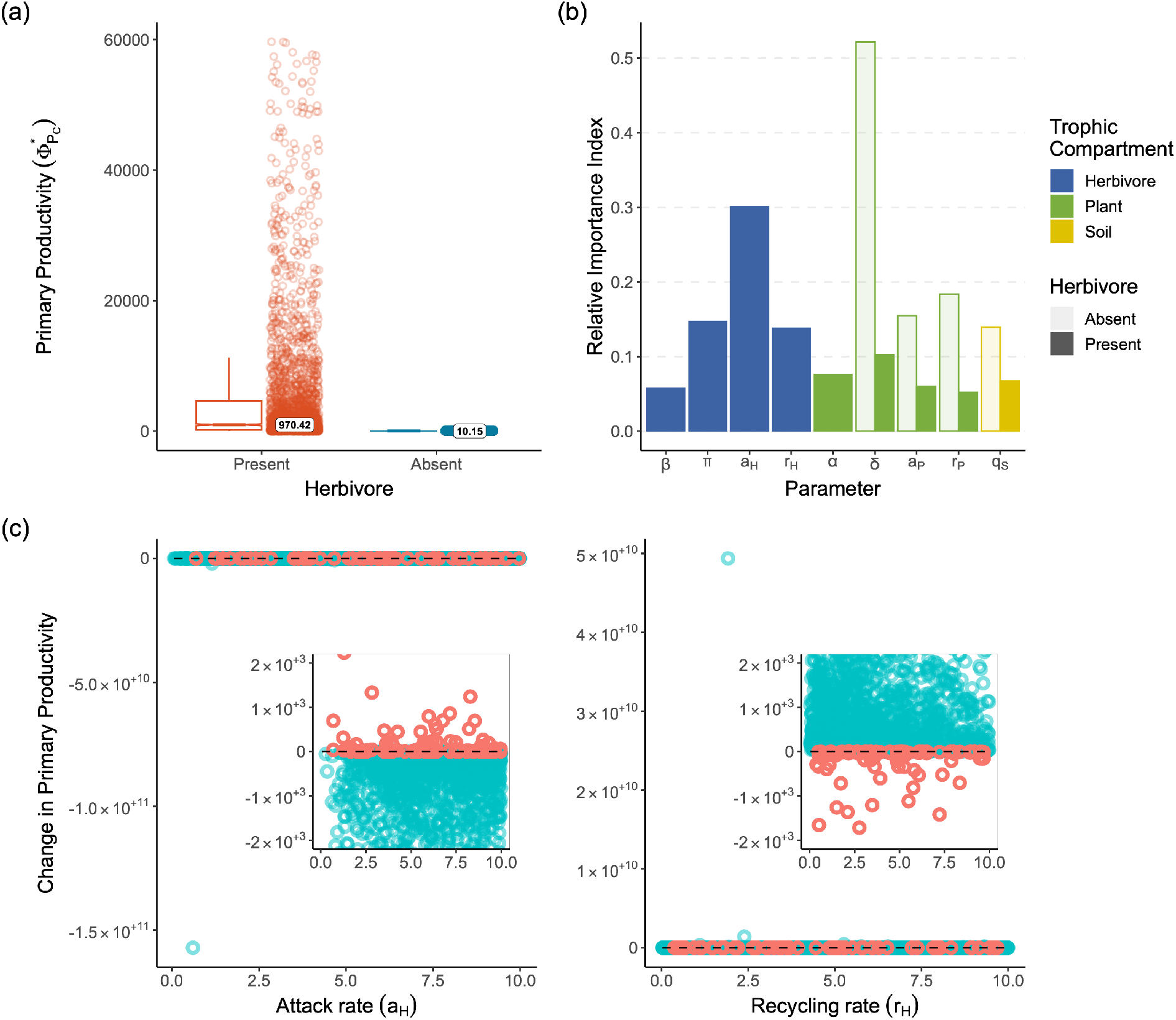
Effects of herbivores’ presence and absence on primary productivity. Panel (a): difference in median primary productivity when herbivores are present vs. when they are absent. The thick line is the median, and the top and bottom hinges are the first and third quantiles. Whiskers extend from the top and bottom hinges to no further than 1.5 times the interquantile range. To avoid graphical artifacts, we removed the top and bottom 5% of data points before drawing the box-and-whisker plot. Panel (b): relative parameter importance as determined using a global stability analysis on the values of primary productivity calculated for carbon. When herbivores are present (solid bars), parameters related to their functional response (𝑎_𝐻_, 𝜋, and 𝑟_𝐻_) are disproportionately more important than any other parameter in the model in determining the primary productivity of the system. In conditions of herbivores’ absence, primary productivity appears driven by parameters pertaining to the functional response of plants (𝛿, 𝑎_𝑃_, 𝑟_𝑃_). Panels (c) and (d): change in primary productivity with increasing herbivore attack rate (𝑎_𝐻_; panel (c)) and recycling rate (𝑟_𝐻_; panel (d)). As expected, increasing values of herbivore attack rate tend to reduce primary productivity whereas higher rates of recycling on part of the herbivore increase it. Red dots identify data points, hence parameter sets, that go counter to these expectations. That is, instances in which increasing values of 𝑎_𝐻_ increase primary productivity and in which increasing 𝑟_𝐻_ decrease it. Insets in both panels zoom into the 𝑦 = 0 area of the graph, which is otherwise difficult to interpret given the wide range of values on the y-axis.

### 4.3 Direct and indirect herbivore effects on Net Ecosystem Carbon Balance

The ecosystem with herbivores had higher estimates of NECB overall compared to the ecosystem without these consumers (Figure 3a). In particular, herbivore presence increased NECB three-fold compared to ecosystem without herbivores (Figure 3a), mirroring the increase in primary productivity observed for the plant compartment (Figure 2a). Again, this increase in NECB appears driven by parameters related to the functional response of the herbivore, in particular the herbivore’s recycling (𝑟_𝐻_) and attack (𝑎_𝐻_) rates (Figure 3b). When herbivores are absent from the system, NECB estimates are driven largely by the primary producers’ uptake rate (𝑎_𝑃_), their recycling rate (𝑟_𝑃_), and the nutrient leaching rate from the soil compartment (𝑞_𝑆_). Conversely, it is harder to identify a single, main driver of NECB in the model with herbivores. While 𝑎_𝑃_, 𝑟_𝑃_, and 𝑞_𝑆_ maintain some weight in shaping NECB in this scenario, herbivore-related parameters appear more influential (Figure 3b)—providing further evidence of the permeating effects of the herbivore-induced trophic rewiring and feedbacks throughout the ecosystem. In particular, the herbivores’ recycling rate (𝑟_𝐻_), attack rate (𝑎_𝐻_), and the plants’ C:N ratio (𝛼) all appear more important for determining NECB in this scenario. The plants’ C:N ratio influence is particularly key, suggesting that NECB is partially influenced by the dietary, quality (C:N)-quantity trade-offs faced by foraging herbivores. Moreover, in the presence of herbivores, plant stoichiometry is relatively more important in shaping NECB than does primary productivity (compare solid bars for 𝛼 between Figure 2b and Figure 3b). While herbivore attack and recycling rates appear to be key in shaping NECB when herbivores are present, we do not observe a clear trend of increase/decrease in NECB with increasing values of either parameter (Figure 3c). Indeed, average estimates of NECB appear higher when herbivore attack rates are low (i.e., herbivore eating little plant biomass) and herbivore recycling is high (i.e., high nutrient returns to the soil via the top-down, “short-circuit” pathway; Figure 1c).

**Figure 3:**
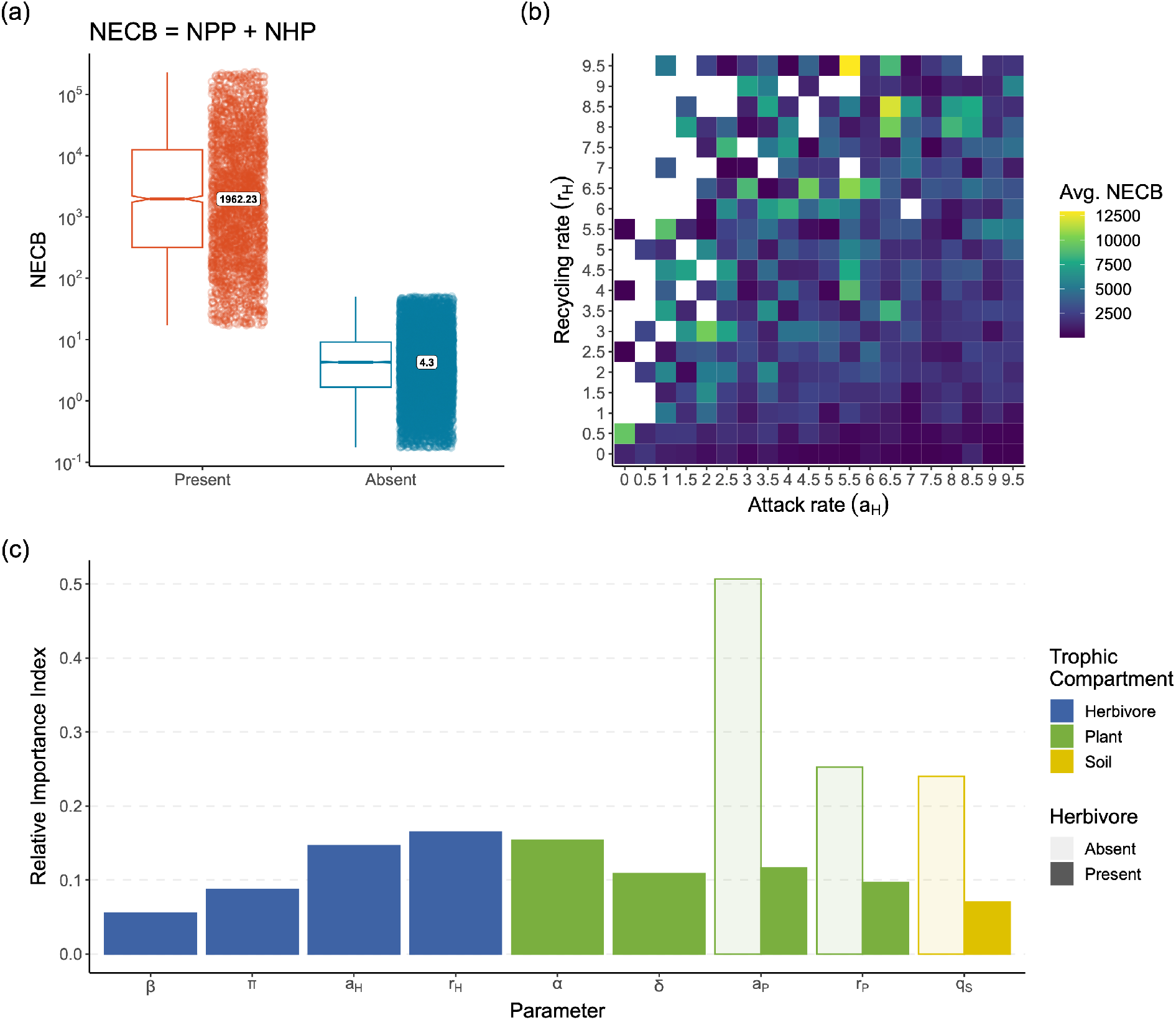
Summary of results of the analyses on Net Ecosystem Carbon Balance (NECB) using equation (4). Panel (a): NECB estimates from the new equation for ecosystem without (blue) and with (red) herbivores. Herbivore presence leads to a three-fold increase in NECB, compared to when ecosystems are devoid of herbivores. Panel (b): heatmap of average NECB estimates for increasing values of herbivore attack (𝑎_𝐻_, x-axis) and recycling (𝑟_𝐻_, y-axis) rates. Panel (c): relative parameter importance in estimating NECB from equation (4) for the model without (light colors) and with (solid colors) herbivores. When herbivores are absent from the system, the plant’s attack rate (𝑎_𝑃_) is the single most important parameter driving NECB estimates. Plant recycling rate (𝑟_𝐻_) also plays a significant, if smaller, role on par with the soil leaching rate (𝑞_𝑆_). Conversely, in ecosystems with herbivores, no clear dominant parameter stands out. Parameters capturing the effects of herbivory—𝑎_𝐻_, 𝑟_𝐻_— show higher values of relative parameter importance, further signalling the strength of the trophic rewiring operated by the consumers’ presence on the functioning of the ecosystem. Plant C:N ratio, 𝛼, appears to play a similarly important role, possibly hinting at dietary trade-offs faced by herbivores. Several other parameters, 𝑎_𝑃_, 𝑟_𝑃_, 𝜋, 𝛿, appear to contribute to shaping NECB estimates when herbivores are present. Notably, soil leaching (𝑞_𝑆_) appears much less relevant for NECB when herbivores are present.

## 5 Discussion

We provide a general framework to account for animal effects on ecosystem carbon cycling and balance. Our framework unites a compartment-style ecosystem model common in community ecology (Daufresne and Loreau 2001a,b; Loreau 2010) to a carbon budget model common in biogeochemistry (Chapin, Matson, and Vitousek 2002; Chapin, Woodwell, et al. 2006), thus laying the foundation for zoogeochemical theory for carbon cycling (*sensu* Schmitz, Wilmers, et al. 2018). Our modeling reveals that ignoring animals when accounting for ecosystem carbon balance could not only result in *quantitatively* inaccurate estimates of ecosystem carbon storage (Figure 2a and 3a), but could also overlook the *qualitatively* different ways in which carbon cycling is controlled when animals are absent compared to when they are present (Figure 1b vs 1d). The latter has important implications for managing ecosystem carbon storage through nature-based climate change solutions that focus merely on plants and soils regardless of the presence of animals.

Herbivore presence restructures the ecosystem’s trophic food web and rewires the cycles of carbon and nitrogen (Figure 1). Consistent with earlier modeling, we found that herbivores cause cascading effects that limit plant growth and foster accumulation of nutrients in soil stocks (e.g., nitrogen in Figure D.2; de Mazancourt, Loreau, and Abbadie 1998; Leroux and Loreau 2010; Loreau 2010). In our modeling these effects further lead to higher storage of atmospheric carbon in plant biomass in the presence of herbivores than in their absence (Figure D.1). This cascading effect influences the balance of C and N stocks in the ecosystem (Figure D.2) by shifting control of cycling from lower trophic compartments to the herbivores (Figure 1b, d). This *qualitatively* different ecosystem control results from the presence of a suite of direct, consumptive effects—captured, for instance, by herbivore consumption (𝑎_𝐻_)—and indirect, non-consumptive processes (i.e., herbivore recycling, 𝑟_𝐻_, and respiration, 𝜋) which supersede their plant-mediated counterparts (parameters 𝑎_𝑃_ and 𝑟_𝑃_; Figure 1d), leading to (i) a short-circuiting of the recycling pathways that replenish the carbon and nitrogen soil pools (DeAngelis 1992) and (ii) the creation of conditions for increased plant nutrient uptake and growth by removing plant biomass (i.e., grazing optimization, Figure 2a; *sensu* Hilbert et al. 1981; de Mazancourt, Loreau, and Abbadie 1998). Herbivore control also partially plays out by limiting the influence of key physical environmental factors (Figure 1d), such as inorganic nutrient inflow (𝐼) or leaching loss (𝑞_𝑆_). Thus, our model’s results offer initial theo-retical support for empirical descriptions of the pervasive influences herbivores can have on an ecosystem’s ability to capture and sequester atmospheric carbon, and on its overall carbon budget (Schmitz, Raymond, et al. 2014; Schmitz, Wilmers, et al. 2018).

The herbivore-caused qualitative changes to the structure of the ecosystem’s trophic food web in turn *quantitatively* alter key ecosystem processes. Estimates for both primary production and net ecosystem carbon balance appear generally higher with herbivores in the ecosystem than without (Figure 2a and 3a, respectively). This increased production and cycling of carbon is consistent with trophic dynamic theory specifying that more productivity is needed to support trophic levels above primary producers (i.e., the ecosystem exploitation hypothesis; Oksanen et al. 1981). However, herbivore influence does not act in only one direction. Primary production appears to respond in contrasting ways to the consumptive and non-consumptive influences exerted by herbivores (Figure 2c, d). As expected, herbivore consumption of plant biomass (parameter 𝑎_𝐻_) leads to a reduction of primary production for most parameter sets (shown in blue in Figure 2c). However, a non-trivial number of parameter sets leads to an increase of primary production when herbivores are present (red dots in Figure 2c). In a similar way, recycling of nutrient from the herbivore compartment (parameter 𝑟_𝐻_) generally increases primary production but a sizeable subset of parameter sets reduces it (blue vs red dots in Figure 2d). The balance of these two processes and the environmental context in which they take place are key for their outcome. Empirical evidence supports these contrasting outcomes (Subalusky and Post 2019; Pringle et al. 2023): while herbivores often limit or reduce primary production (Tanentzap and Coomes 2012), under certain conditions they can enhance it—for instance, as do the hippopotamuses of the Mara River (Subalusky, Dutton, Rosi-Marshall, et al. 2015; Subalusky, Dutton, Njoroge, et al. 2018) or the ungulate assemblages in several grassland ecosystems in Africa, Eurasia, and North America (McNaughton 1979; Frank and McNaughton 1993; Schoenecker, Zeigenfuss, and Augustine 2022).

Likewise, it is difficult to identify a single directional effect of herbivore consumption and recycling on NECB. Herbivore presence predominantly increases NECB (Figure 3a) and partially limits plant control over this process (compare light and solid bars in Figure 3b). However, varying levels of herbivore consumption (𝑎_𝐻_) and recycling (𝑟_𝐻_) can have varying effects on NECB estimates (Figure 3c) which depend on specific parameter combinations. This outcome is consistent with empirical evidence that herbivores can enhance or limit carbon capture but the effects are highly case-study-specific (see review in Pringle et al. 2023). Even species that are phylogenetically similar can have different effects on carbon dynamics. Forest elephants (*Loxodonta cyclotis*) in the rainforests of the Congo Basin, in central Africa, enhance carbon storage by primary producers through foraging effects and other types of indirect disturbance (e.g., debarking, trampling; Berzaghi, Bretagnolle, et al. 2023). Elsewhere in Africa, high densities of bull savanna elephants (*L. africana*) lead to reduction in aboveground carbon gains in the subtropical savanna of Kruger National Park (Davies and Asner 2019).

Together with this growing body of empirical evidence (see Schmitz, Wilmers, et al. 2018; Schmitz, Sylvén, et al. 2023; Pringle et al. 2023), our model shows that any effort to accurately account for the uptake, cycling, and storage of carbon in ecosystems should include animal contributions. Simply implicitly accounting for animal-mediated processes via heterotrophic respiration does not return a complete picture of an ecosystem’s carbon balance and indeed appears to under-estimate it—as shown by the results of our model when herbivores are absent (in blue in Figures 2 and 3; see also Rastetter et al. 2022). Empirical (Holdo et al. 2009; Malhi, Doughty, et al. 2016; Doughty et al. 2016; Don et al. 2019; Berzaghi, Longo, et al. 2019; Leroux, Wiersma, and Vander Wal 2020; Sitters et al. 2020), and now theoretical, evidence increasingly challenges the assumption that an ecosystem’s biomass-dominant compartments (plant, microbes) can subsume the effects of animals on the carbon cycle.

Beyond the risk of inaccurately estimating an ecosystem’s carbon budget, the model results have important implications for any management effort aimed at enhancing ecosystem carbon storage using conventional means. For example, simply increasing primary producers’ biomass via planting and cultivation—would be using the wrong lever to achieve its goals. Instead, and perhaps counterintuitively, protecting and building the biomass of animal populations can instigate feedback loops—through the consumptive and recycling pathways highlighted above—that can influence and potentially enhance an ecosystem’s primary production and net carbon balance. While much remains to be clarified (Schmitz, Wilmers, et al. 2018; Kristensen et al. 2022; Pringle et al. 2023), theoretical and empirical evidence increasingly supports reorienting our efforts to develop nature-based climate change solutions that explicitly incorporate animals (Berzaghi, Cosimano, et al. 2022; Malhi, Lander, et al. 2022; Schmitz, Sylvén, et al. 2023; Schmitz and Sylvén 2023). We offer our model as a scaffolding to provide general insights on the role of animals in ecosys-tem carbon cycling and balance. The exact amounts of carbon captured and stored in an ecosystem will depend on the assemblage of animal, plant, and microbial species comprising a given ecosystem and on its biogeophysical conditions—e.g., soil composition, temperature, moisture, etc. The interplay between these factors will have a decided effect on mechanisms and rates of carbon uptake via photosynthetic capture and assimilation, consumptive transfer and accumulation in animal (herbivore and carnivore) biomass, microbial processes, and on the rates of autotrophic (plant) and heterotrophic (microbial and animal) release of carbon into the atmosphere. It follows that applying the modeling framework we describe here to a particular ecosystem will require gathering and substituting model parameter values for functions which explicitly incorporate physiological mechanisms and whose values vary across the domain of biophysical conditions present in the ecosystem, as is done with conventional carbon cycle models that exclude animals (Piao et al. 2013; Zaehle et al. 2014) and with animal-driven models in certain ecosystems (Dangal et al. 2017; Rastetter et al. 2022). Indeed, empirical knowledge to quantify animal-mediated processes and thus parameterize the model for specific systems already exists for several ecosystems and animal species (reviewed in Pringle et al. 2023) and could allow our model to inform restoration and management efforts aimed at leveraging animal contributions to increase ecosystem carbon capture and storage (e.g., through trophic rewilding; *sensu* Schmitz, Sylvén, et al. 2023; Schmitz and Sylvén 2023).

We present a simple model rooted in the trophic compartment formalization—consistent with previous modeling efforts. This approach implicitly assumes that the many species comprising a trophic compartment are all functionally equivalent (Schmitz and Leroux 2020). That is, the model does not differentiate between a hare and an elephant as both can be described as “herbivores”. Yet, ecosystems contain diverse assemblages of species that may share some characteristics (e.g., diet type) but not others thus need not be assumed as functionally equivalent (Schmitz and Leroux 2020). While helpful in reducing the mathematical complexity of the model and maintaining its analytical tractability, assuming functional equivalence limits our ability to disentangle the role of individual species’ traits in enabling their influence on ecosystem processes. As well, and consequently, it also constraints the usefulness of this type of model in exploring the relationship between ecosystem functioning and animal biodiversity. A possible way to address this limitation and dissect functional differences among multiple species within a trophic compartment would be to expand our model framework to incorporate functional traits of animals—i.e., the morphological (e.g., bird beak shape), behavioral (e.g., group vs solitary living), or physiological (e.g., ruminant vs non-ruminant) attributes that drive an organism’s biogeochemical function (*sensu* Schmitz and Leroux 2020). Our model uses a linear (type I) functional response to represent both plant uptake of nutrients from the soil and herbivore consumption of plant biomass, consistent with earlier ecosystem models of elemental flux among trophic compartments. Assuming a type I functional response allows for simpler model formulation, enhanced mathematical tractability, and comparison to other models (Jonsson 2017). However, more complex functional responses may be worth exploring to develop a more complete picture of animal control on carbon cycling. Finally, our framework does not explicitly model microbial activities connected to the carbon cycle—respiration and production—subsuming them into the soil compartment (Figure 1a, c). Dissecting the organic and inorganic components of soil—for instance, by adding a detritus and decomposer trophic compartments to the model—would shed light on the interactions between the above- and below-ground food webs, their individual and combined effects on ecosystem carbon cycling, and on their synergistic or antagonistic nature (Kristensen et al. 2022; Naidu, Roy, and Bagchi 2022; Losada et al. 2023).

Carbon capture, cycling, and storage are essential ecosystem processes to help arrest climate change. Historically, models used to quantify these processes have excluded animals from their frameworks by reasoning that animals make up a small portion of an ecosystem’s total biomass and this limits their influence of carbon dynamics. Increasingly, empirical evidence shows that this is not the case and rather animal influences permeate all aspects of ecosystem carbon budgets. Here, we develop an ecosystem carbon model that accounts for animal control of ecosystem carbon dynamics by focusing on herbivores. We show that (i) herbivore presence rewires the ecosystem’s trophic food web and qualitatively changes how its carbon cycle plays out, concentrating control in the top trophic compartment; (ii) this re-arrangement of the carbon cycle leads to quantitative changes in key ecosystem processes involved in the ecosystem’s carbon budget; and (iii) generalizing herbivore effects across different conditions and scenarios can be challenging as herbivores’ control and influence on ecosystem carbon cycle varies with context. Together, these results lend further support to calls for integrating estimates of animal influences on ecosystem carbon services in existing models of carbon cycling at multiple spatio-temporal scales and to the development of nature-based solutions that leverage animal contributions to mitigate climate change, such as, trophic rewilding.

## Data Availability

All data and code used in our analyses is available through the online repository at: https://doi.org/10.6084/m9.figshare.23688855.

## Appendices

In these appendices, we provide additional details on the derivation and analyses of both the herbivore and non-herbivore models discussed in the main text. In Appendix A, we show additional equilibria obtained for both models. Appendix B shows the derivation of function 𝑊_𝐶𝐻_[𝐻_𝑁_, 𝐻_𝐶_] as well as show the relationship between parameters 𝛼 and 𝛽. In Appendix C, we present the derivation of the NECB equations used to calculate the values of this ecosystem function in both modeling scenarios—*with* and *without* herbivores—and for both NECB formulae used. Finally, in Appendix D we present additional figures where we report results on the effects of herbivore presence and absence on equilibrium soil stocks and plant biomass.

### A Model equilibria and feasibility conditions

#### A.1 Model without Herbivores

##### Equilibria

The model without herbivores has two equilibria: one where all state variables values are positive and thus biologically meaningful (i.e., “feasible” equilibrium), and one where they are not. The solutions for these equilibria are:

Feasible Equilibrium

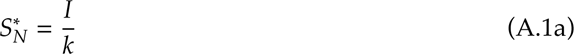

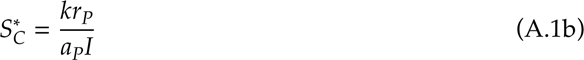

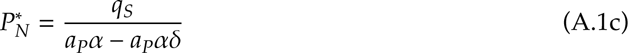

Non-feasible equilibrium

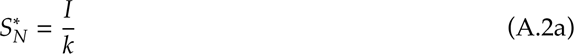

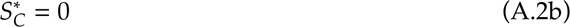

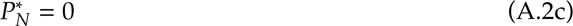

##### Feasibility conditions

In the feasible equilibrium, all state variables are > 0 when 𝛼 > 0, 0 < 𝛿 < 1, 𝑟_𝑃_ > 0, 𝑞_𝑆_ > 0, 𝑘 > 0, 𝐼 > 0, and 𝑎_𝑃_ > 0.

#### A.2 Model with Herbivores

##### Equilibria

The model with herbivores has three equilibria: one where are all state variables are > 0 and thus biologically meaningful, and two where they are not.

Feasible equilibrium:

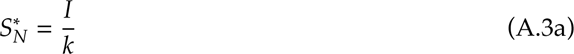

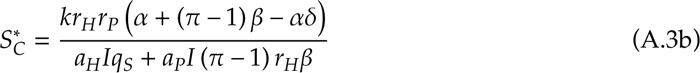

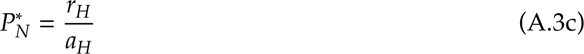

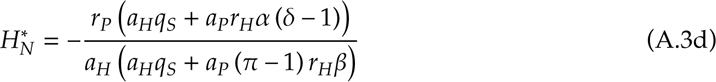

The two non-feasible equilibria are,

Case 1:

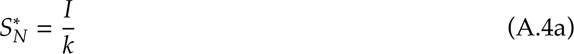

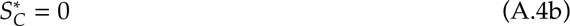

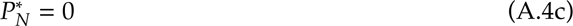

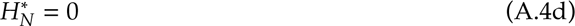

Case 2:

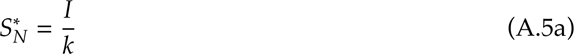

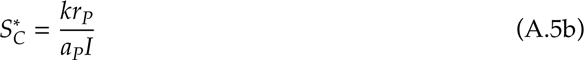

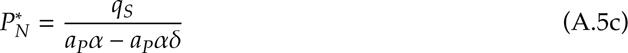

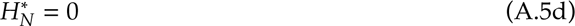

##### Feasibility conditions

The feasible equilibrium produces positive stocks for all state variables when 𝛼 > 0, 𝛽 > 0, 𝑎_𝐻_ > 0, 𝑎_𝑃_ > 0, 𝐼 > 0, 𝑘 > 0, 𝑟_𝐻_ > 0, 𝑟_𝑃_ > 0, and

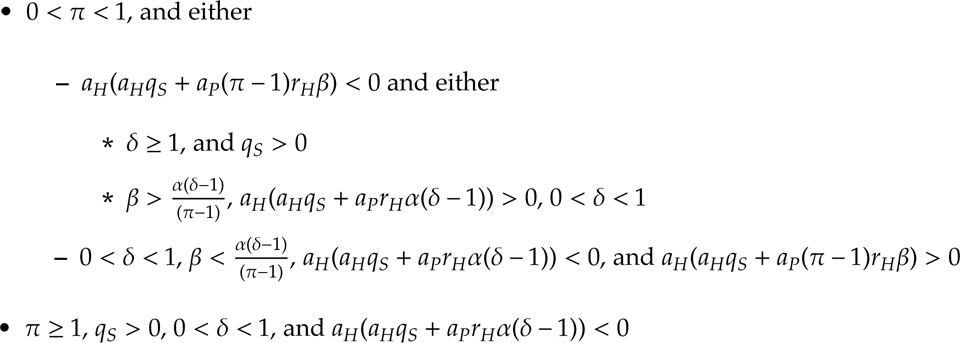

#### B Derivation of herbivore differential assimilation rates

Here, we derive the expressions for the term 𝑊_𝐶𝐻_[𝐻_𝑁_, 𝐻_𝐶_] in the model (Table 1), the fraction of assimilated C that herbivore release over time as a function of their content of C and N.

##### B.1 Herbivore differential assimilation rate for Carbon

In our model, herbivores are N-limited; that is, herbivores eliminate excess C obtained through diet from their system via respiration. Note that, in our model, equation (2f) and equation (2e) are related through parameter 𝛽—the herbivore’s C:N ratio—according to the following equation:

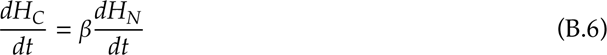

It follows that,

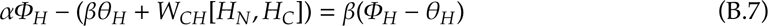

Expanding the term in parentheses on the left hand side of the equation, and isolating the 𝑊_𝐶𝐻_[𝐻_𝑁_, 𝐻_𝐶_], we get:

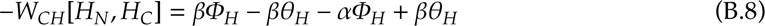

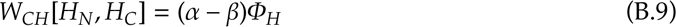

And, finally, substituting the formula for 𝛷_𝐻_ (Table 1) in equation (B.9), we get the formula for 𝑊_𝐶𝐻_[𝐻_𝑁_, 𝐻_𝐶_] as shown in Table 1:

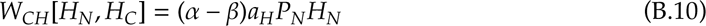

#### C Net Ecosystem Carbon Balance Derivation

##### Model without Herbivores

For the model *without* herbivores, NPP is

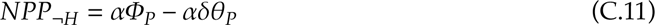

Likewise, NHP for the model *without* herbivores comprises Soil-related terms only,

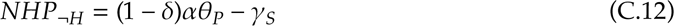

Substituting Equations (C.11) and (C.12) into Equation (4), we get

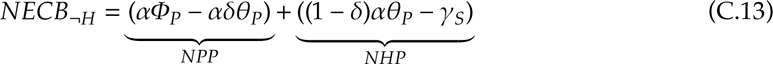

Expanding the products, we get the equation for NECB *without* herbivores,

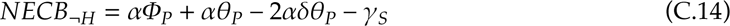

##### Model with Herbivores

Both NPP and NHP are more complex due the presence of additional parameters—namely, 𝛼 and 𝛽—that represent the C:N ratio of plants and herbivores, respectively. NPP has the formula

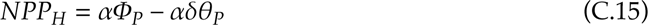

For NHP, the formula is

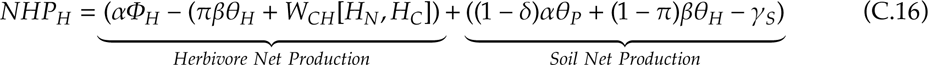

Expanding the (1−𝜋)𝛽𝜃_𝐻_ product in the Soil Net Production, we can cancel out the 𝜋𝛽𝜃_𝐻_ terms,

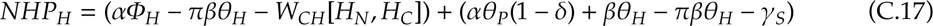

And then, plugging in the value of 𝑊_𝐶𝐻_[𝐻_𝑁_, 𝐻_𝐶_] from Table 1,

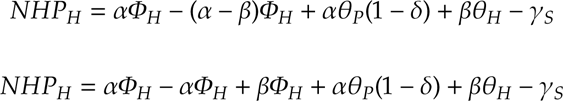

So that the NHP equation *with* herbivores is

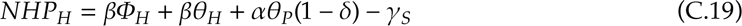

Now, we can substitute the values of NPP (Equation (C.15)) and NHP (Equation (C.19)) in the equation for NECB (Equation (4))

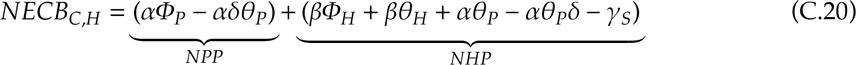

Notice that we expanded the 𝛼𝜃_𝑃_(1 − 𝛿) product in Equation (C.20). As the 𝛼𝛿𝜃_𝑃_ terms sum up, Equation (C.20) simplifies to

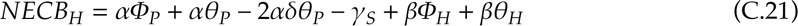

Substituting in Equation (C.21) the formulae for 𝛷_𝑃_, 𝛷_𝐻_, 𝜃_𝑃_, 𝜃_𝐻_, and 𝛾_𝑆_ from Table 1 and the equilibrium values of our state variables—𝑆^∗^_𝐶_, 𝑃^∗^_𝐶_, 𝐻_𝐶_^∗^ —produces the equation to use in our analyses.

As expected, Equation (C.21) resembles Equation (C.14) with the terms that relate to the herbivores.

#### D Additional figures

**Figure D.1:**
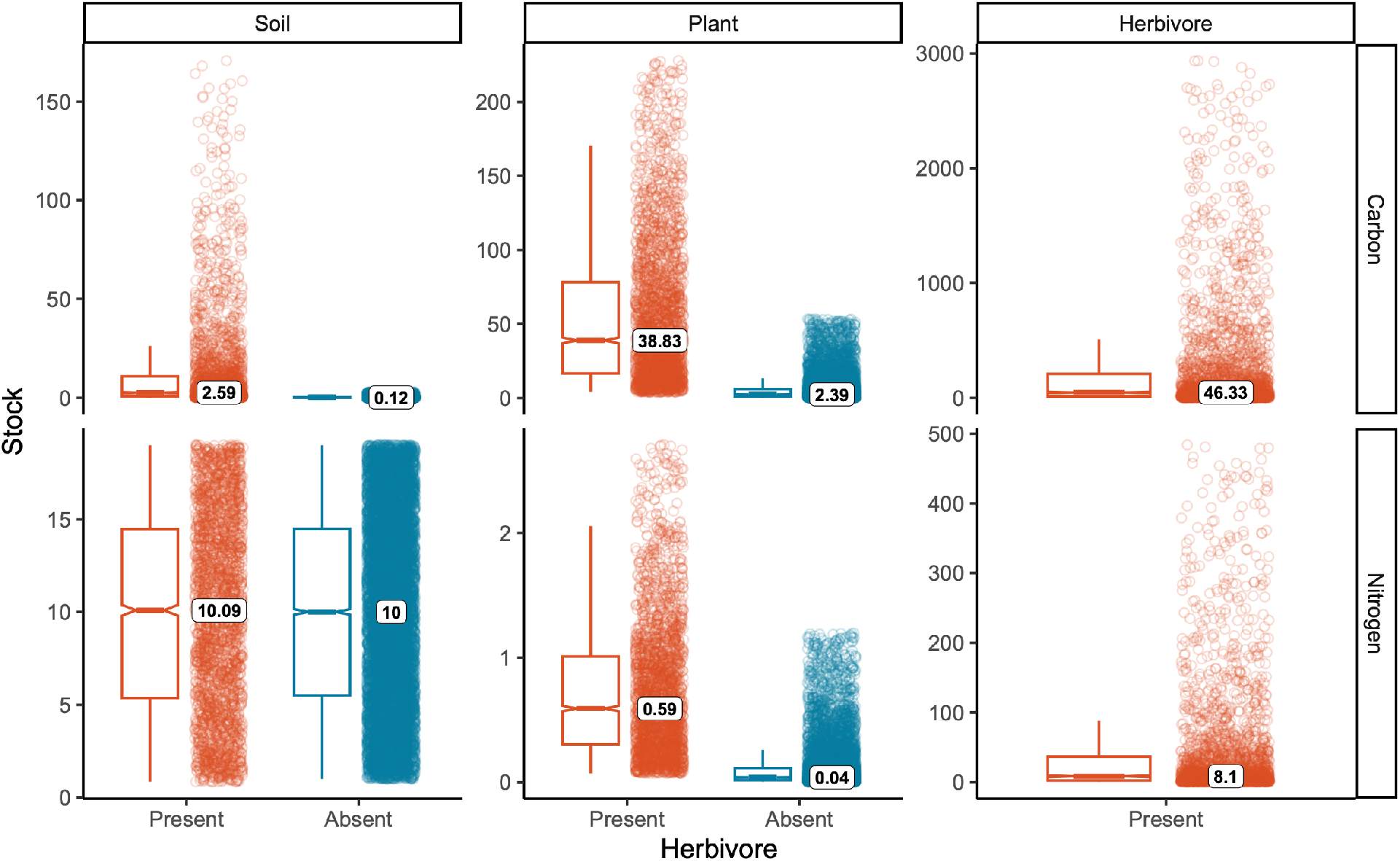
Nutrient content for each trophic compartment in the ecosystem at equilibrium, with and without herbivores. At equilibrium, herbivore presence (red box-and-whiskers plots, jitter) leads to a one-fold increase in C content (top row) in both soil and plant trophic compartments compared to the model *without* herbivores (blue box-and-whiskers plots, jitter). Nitrogen is found mostly in the soil stock when herbivores are absent from the system, likely as a consequence of the strong bottom-up control present in this ecosystem (Figure 1b). Herbivore presence allows for a trophic cascade to emerge, with N being equally abundant in soil and herbivores and almost entirely absent from plant biomass—something that is more clearly visibly when standardizing these amounts by the total biomass in the system (Figure D.2). Note the different scales of both x- and y-axes in all six panels.

**Figure D.2:**
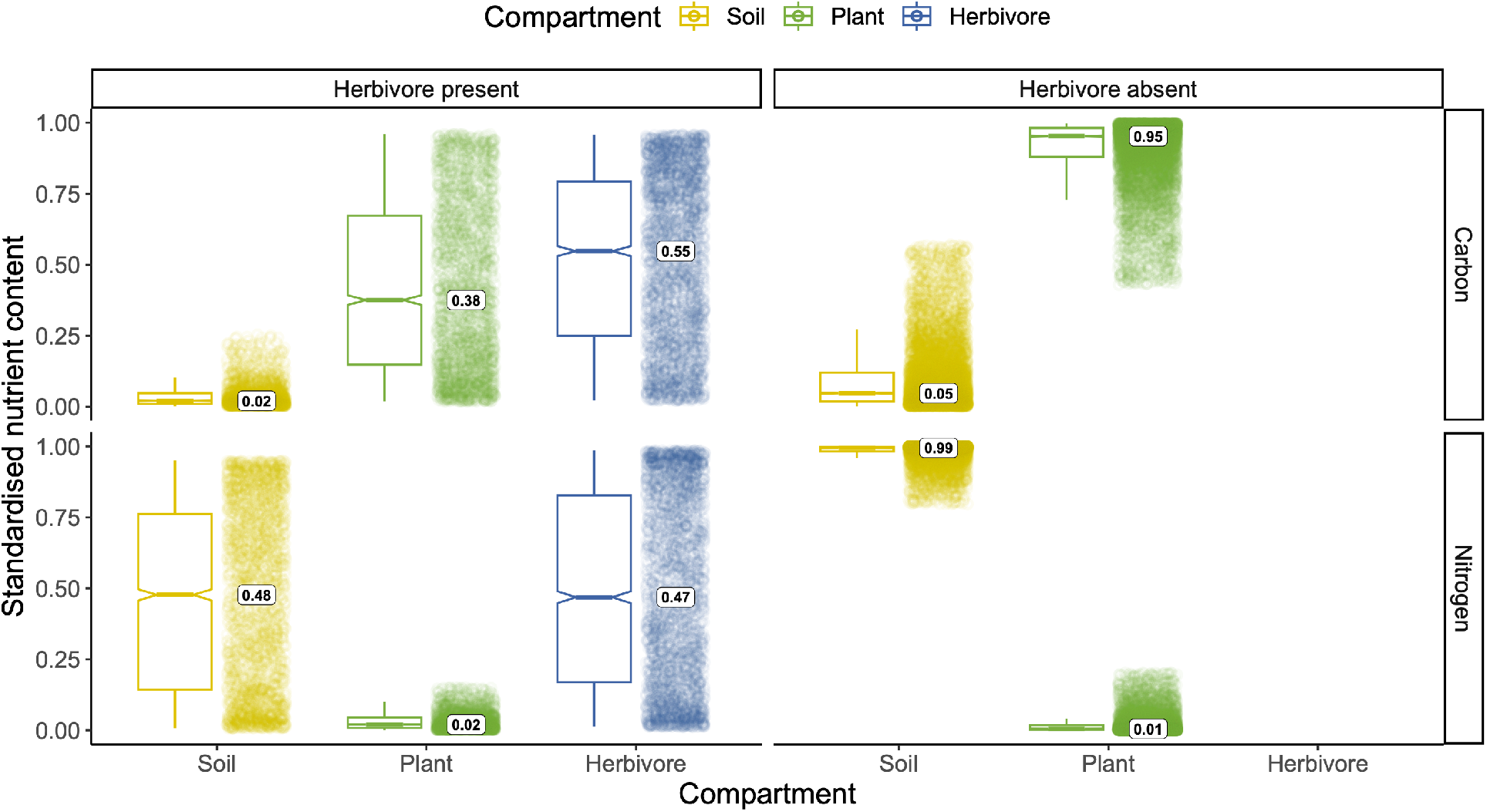
Standardized nutrient content for each trophic compartment in the ecosystem at equilibrium, with and without herbivores. Carbon (C) nutrient content (top row) appears to accumulate in higher trophic compartments. However, while it appears a larger share of C is locked in plant biomass when herbivores are absent from the ecosystem, in absolute terms this is a much smaller amount than when herbivores are present (Figure D.1). Conversely, nitrogen (N) appears to strictly adhere to top-down, trophic-cascade-like dynamics when herbivores are present and to bottom-up dynamics when they are absent (bottom row). Again, while it would appear that more N is capture by the ecosystem in the absence of herbivores, in absolute terms this is a much larger quantity (Figure D.1). Standardized nutrient content values were obtained by dividing each trophic compartment’s C or N content by the total C or N in the ecosystem, for either scenario.

## References

Ballantyne, A. P. et al. (2021). “Reconciling carbon-cycle processes from ecosystem to global scales”. Frontiers in Ecology and the Environment 19.1, pp. 57–65. DOI: 10.1002/fee.2296.

Bassar, R. D. et al. (2012). “Direct and Indirect Ecosystem Effects of Evolutionary Adaptation in the Trinidadian Guppy (Poecilia reticulata).” The American Naturalist 180.2, pp. 167–185. DOI: 10.1086/666611.

Bellmore, J. R. et al. (2014). “The response of stream periphyton to Pacific salmon: using a model to understand the role of environmental context”. Freshwater Biology 59.7, pp. 1437–1451. DOI: 10.1111/fwb.12356.

Berzaghi, F., F. Bretagnolle, et al. (2023). “Megaherbivores modify forest structure and increase carbon stocks through multiple pathways”. Proceedings of the National Academy of Sciences 120.5. Publisher: Proceedings of the National Academy of Sciences, e2201832120. DOI: 10.1073/pnas.2201832120.

Berzaghi, F., T. Cosimano, et al. (2022). “Value wild animals’ carbon services to fill the biodiversity financing gap”. Nature Climate Change 12.7, pp. 598–601. DOI: 10.1038/s41558-022-01407-4.

Berzaghi, F., M. Longo, et al. (2019). “Carbon stocks in central African forests enhanced by elephant disturbance”. Nature Geoscience 12.9, pp. 725–729. DOI: 10.1038/s41561-019-0395-6.

Buchkowski, R. W., S. J. Leroux, and O. J. Schmitz (2019). “Microbial and animal nutrient limitation change the distribution of nitrogen within coupled green and brown food chains”. Ecology 100.5, e02674. DOI: 10.1002/ecy.2674.

Carnell, R. (2022). lhs: Latin Hypercube Samples. R package version 1.1.5. URL: https://CRAN.R-project.org/package=lhs.

Chapin, F. S., P. A. Matson, and P. M. Vitousek (2002). Principles of Terrestrial Ecosystem Ecology. 2nd ed. New York, NY: Springer New York, NY. URL: https://doi.org/10.1007/978-1-4419-9504-9.

Chapin, F. S., G. M. Woodwell, et al. (2006). “Reconciling carbon-cycle concepts, terminology, and methods”. Ecosystems 9.7, pp. 1041–1050. DOI: 10.1007/s10021-005-0105-7.

Dangal, S. R. S. et al. (2017). “Integrating Herbivore Population Dynamics Into a Global Land Biosphere Model: Plugging Animals Into the Earth System”. Journal of Advances in Modeling Earth Systems 9.8, pp. 2920–2945. DOI: 10.1002/2016MS000904.

Daufresne, T. and M. Loreau (2001a). “ECOLOGICAL STOICHIOMETRY, PRIMARY PRODUCER– DECOMPOSER INTERACTIONS, AND ECOSYSTEM PERSISTENCE”. Ecology 82.11, pp. 3069– 3082. DOI: 10.1890/0012-9658(2001)082[3069:ESPPDI]2.0.CO;2.

Daufresne, T. and M. Loreau (2001b). “Plant-herbivore interactions and ecological stoichiometry: when do herbivores determine plant nutrient limitation?” Ecology Letters 4.3, pp. 196–206. DOI: 10.1046/j.1461-0248.2001.00210.x.

Davies, A. B. and G. P. Asner (2019). “Elephants limit aboveground carbon gains in African savannas”. Global Change Biology 25.4, pp. 1368–1382. DOI: 10.1111/gcb.14585.

de Mazancourt, C., M. Loreau, and L. Abbadie (1998). “Grazing Optimization and Nutrient Cycling: When Do Herbivores Enhance Plant Production?” Ecology 79.7, pp. 2242–2252. DOI: 10.2307/176819.

DeAngelis, D. L. (1992). Dynamics of nutrient cycling and food webs. Ed. by J. W. Shipley. Vol. 9. Population and Community Biology Series. Springer Dordrecht, p. 288. DOI: 10.1007/978-94-011-2342-6.

Don, A. et al. (2019). “Simulated wild boar bioturbation increases the stability of forest soil carbon”. Biogeosciences 16.21, pp. 4145–4155. DOI: 10.5194/bg-16-4145-2019.

Doughty, C. E. et al. (2016). “Megafauna extinction, tree species range reduction, and carbon storage in Amazonian forests”. Ecography 39.2, pp. 194–203. DOI: 10.1111/ecog.01587.

Frank, D. A. and S. J. McNaughton (1993). “Evidence for the promotion of aboveground grassland production by native large herbivores in Yellowstone National Park”. Oecologia 96.2, pp. 157–161. DOI: 10.1007/BF00317727.

Gravel, D. et al. (2010). “Source and sink dynamics in meta-ecosystems”. Ecology 91.7, pp. 2172– 2184. DOI: 10.1890/09-0843.1.

Hall, S. R. et al. (2007). “Food quality, nutrient limitation of secondary production, and the strength of trophic cascades”. Oikos 116.7, pp. 1128–1143. DOI: 10.1111/j.2007.0030-1299.15875.x.

Harper, E. B., J. C. Stella, and A. K. Fremier (2011). “Global sensitivity analysis for complex ecological models: a case study of riparian cottonwood population dynamics”. Ecological Applications 21.4, pp. 1225–1240. DOI: 10.1890/10-0506.1.

Hilbert, D. W. et al. (1981). “Relative growth rates and the grazing optimization hypothesis”. Oecologia 51.1, pp. 14–18. DOI: 10.1007/BF00344645.

Holdo, R. M. et al. (2009). “A Disease-Mediated Trophic Cascade in the Serengeti and its Implications for Ecosystem C”. PLOS Biology 7.9. Publisher: Public Library of Science, e1000210. DOI: 10.1371/journal.pbio.1000210.

Holt, R. D. (1997). “Community modules”. In: Multitrophic Interactions in Terrestrial Ecosystems. A.C. Gange and V.K. Brown, eds. Hoboken, NJ: Blackwell Science, pp. 333–349. URL: https://people.clas.ufl.edu/rdholt/files/076.pdf.

Huntley, M. E., M. D. Lopez, and D. M. Karl (1991). “Top Predators in the Southern Ocean: A Major Leak in the Biological Carbon Pump”. Science 253.5015, pp. 64–66. DOI: 10.1126/science.1905841.

Ives, A. R., B. J. Cardinale, and W. E. Snyder (2005). “A synthesis of subdisciplines: predator–prey interactions, and biodiversity and ecosystem functioning”. Ecology Letters 8.1, pp. 102–116. DOI: 10.1111/j.1461-0248.2004.00698.x.

Jonsson, T. (2017). “Conditions for Eltonian Pyramids in Lotka-Volterra Food Chains”. Scientific Reports 7.1, p. 10912. DOI: 10.1038/s41598-017-11204-1.

Keenan, T. and C. Williams (2018). “The Terrestrial Carbon Sink”. Annual Review of Environment and Resources 43.1, pp. 219–243. DOI: 10.1146/annurev-environ-102017-030204.

Kristensen, J. A. et al. (2022). “Can large herbivores enhance ecosystem carbon persistence?” Trends in Ecology & Evolution 37.2, pp. 117–128. DOI: 10.1016/j.tree.2021.09.006.

Leroux, S. J., D. Hawlena, and O. J. Schmitz (2012). “Predation risk, stoichiometric plasticity and ecosystem elemental cycling”. Proc. R. Soc. B Biol. Sci. 279.1745, pp. 4183–4191. DOI: 10.1098/rspb.2012.1315.

Leroux, S. J. and M. Loreau (2008). “Subsidy hypothesis and strength of trophic cascades across ecosystems”. Ecology Letters 11.11, pp. 1147–1156. DOI: 10.1111/j.1461-0248.2008.01235.x.

Leroux, S. J. and M. Loreau (2010). “Consumer-mediated recycling and cascading trophic interactions”. Ecology 91.7, pp. 2162–2171. DOI: 10.1890/09-0133.1.

Leroux, S. J. and O. J. Schmitz (2015). “Predator-driven elemental cycling: the impact of predation and risk effects on ecosystem stoichiometry”. Ecology and Evolution 5.21, pp. 4976–4988. DOI: 10.1002/ece3.1760.

Leroux, S. J., Y. F. Wiersma, and E. Vander Wal (2020). “Herbivore Impacts on Carbon Cycling in Boreal Forests”. Trends in Ecology & Evolution 35.11, pp. 1001–1010. DOI: 10.1016/j.tree.2020.07.009.

Loreau, M. (2010). From populations to ecosystems. Ed. by S. A. Levin and H. S. Horn. Monographs in Population Biology. Princeton University Press. DOI: 10.1515/9781400834167.

Losada, M. et al. (2023). “Mammal and tree diversity accumulate different types of soil organic matter in the northern Amazon”. iScience 26.3, p. 106088. DOI: 10.1016/j.isci.2023.106088.

Malhi, Y., C. E. Doughty, et al. (2016). “Megafauna and ecosystem function from the Pleistocene to the Anthropocene”. Proceedings of the National Academy of Sciences 113.4. Publisher: Proceedings of the National Academy of Sciences, pp. 838–846. DOI: 10.1073/pnas.1502540113.

Malhi, Y., T. Lander, et al. (2022). “The role of large wild animals in climate change mitigation and adaptation”. Current Biology 32.4, R181–R196. DOI: 10.1016/j.cub.2022.01.041.

McNaughton, S. J. (1979). “Grazing as an Optimization Process: Grass-Ungulate Relationships in the Serengeti”. The American Naturalist 113.5. Publisher: The University of Chicago Press, pp. 691–703. DOI: 10.1086/283426.

Metcalfe, D. B. et al. (2014). “Herbivory makes major contributions to ecosystem carbon and nutrient cycling in tropical forests”. Ecology Letters 17.3. Ed. by W. van der Putten, pp. 324–332. DOI: 10.1111/ele.12233.

Meunier, C. L. et al. (2017). “From Elements to Function: Toward Unifying Ecological Stoichiometry and Trait-Based Ecology”. Frontiers in Environmental Sciences 5, pp. 1–10. DOI: 10.3389/fenvs.2017.00018.

Middelburg, J. J. (2019). *Marine carbon biogeochemistry: A primer for earth system scientists*. Ed. by G. Lohmann et al. SpringerBriefs in Earth System Sciences. Cham, Switzerland: Springer Nature. DOI: 10.1007/978-3-030-10822-9.

Naidu, D. G. T., S. Roy, and S. Bagchi (2022). “Loss of grazing by large mammalian herbivores can destabilize the soil carbon pool”. Proceedings of the National Academy of Sciences 119.43, e2211317119. DOI: 10.1073/pnas.2211317119.

Oksanen, L. et al. (1981). “Exploitation Ecosystems in Gradients of Primary Productivity”. The American Naturalist 118.2, pp. 240–261. DOI: 10.1086/283817.

Piao, S. et al. (2013). “Evaluation of terrestrial carbon cycle models for their response to climate variability and to CO_2_ trends”. Global Change Biology 19.7, pp. 2117–2132. DOI: 10.1111/gcb.12187.

Pringle, R. M. et al. (2023). “Impacts of large herbivores on terrestrial ecosystems”. Current Biology 33.11, R584–R610. DOI: 10.1016/j.cub.2023.04.024.

R Core Team (2022). R: A Language and Environment for Statistical Computing. R Foundation for Statistical Computing. Vienna, Austria. URL: https://www.R-project.org/.

Rastetter, E. B. et al. (2022). “Model responses to CO_2_ and warming are underestimated without explicit representation of Arctic small-mammal grazing”. Ecological Applications 32.1. DOI: 10.1002/eap.2478.

Rizzuto, M., S. J. Leroux, and O. J. Schmitz (2023). “Code and data for Rewiring the carbon cycle: a theoretical framework for animal-driven ecosystem carbon sequestration.” DOI: 10.6084/m9.figshare.23688855.

Schmitz, O. J. and S. J. Leroux (2020). “Food webs and ecosystems: linking species interactions to the carbon cycle”. Annual Review of Ecology, Evolution, and Systematics 51.1, pp. 271–295. DOI: 10.1146/annurev-ecolsys-011720-104730.

Schmitz, O. J., P. A. Raymond, et al. (2014). “Animating the carbon cycle”. Ecosystems 17.2, pp. 344–359. DOI: 10.1007/s10021-013-9715-7.

Schmitz, O. J. and M. Sylvén (2023). “Animating the Carbon Cycle: How Wildlife Conservation Can Be a Key to Mitigate Climate Change”. Environment: Science and Policy for Sustainable Development 65.3, pp. 5–17. DOI: 10.1080/00139157.2023.2180269.

Schmitz, O. J., M. Sylvén, et al. (2023). “Trophic rewilding can expand natural climate solutions”. Nature Climate Change, pp. 1–10. DOI: 10.1038/s41558-023-01631-6.

Schmitz, O. J., C. C. Wilmers, et al. (2018). “Animals and the zoogeochemistry of the carbon cycle”. Science 362.6419, eaar3213. DOI: 10.1126/science.aar3213.

Schoenecker, K. A., L. C. Zeigenfuss, and D. J. Augustine (2022). “Can grazing by elk and bison stimulate herbaceous plant productivity in semiarid ecosystems?” Ecosphere 13.4, e4025. DOI: 10.1002/ecs2.4025.

Sitters, J. et al. (2020). “Negative effects of cattle on soil carbon and nutrient pools reversed by megaherbivores”. Nature Sustainability 3.5, pp. 360–366. DOI: 10.1038/s41893-020-0490-0.

Steinberg, D. K. and M. R. Landry (2017). “Zooplankton and the Ocean Carbon Cycle”. Annual Review of Marine Science 9.1, pp. 413–444. DOI: 10.1146/annurev-marine-010814-015924.

Sterner, R. W. and J. J. Elser (2002). Ecological stoichiometry: the biology of elements from molecules to the biosphere. Princeton University Press.

Subalusky, A. L., C. L. Dutton, L. Njoroge, et al. (2018). “Organic matter and nutrient inputs from large wildlife influence ecosystem function in the Mara River, Africa”. Ecology 99.11, pp. 2558– 2574. DOI: 10.1002/ecy.2509.

Subalusky, A. L., C. L. Dutton, E. J. Rosi-Marshall, et al. (2015). “The hippopotamus conveyor belt: Vectors of carbon and nutrients from terrestrial grasslands to aquatic systems in sub-Saharan Africa”. Freshwater Biology 60.3, pp. 512–525. DOI: 10.1111/fwb.12474.

Subalusky, A. L. and D. M. Post (2019). “Context dependency of animal resource subsidies”. Biol. Rev. Camb. Philos. Soc. 94.2. DOI: 10.1111/brv.12465.

Tanentzap, A. J. and D. A. Coomes (2012). “Carbon storage in terrestrial ecosystems: do browsing and grazing herbivores matter?” Biological Reviews 87.1, pp. 72–94. DOI: 10.1111/j.1469-185X.2011.00185.x.

Welti, N. et al. (2017). “Bridging food webs, ecosystem metabolism, and biogeochemistry using ecological stoichiometry theory”. Frontiers in Microbiology 8, pp. 1–14. DOI: 10.3389/fmicb.2017.01298.

White, J. W. et al. (2014). “Ecologists should not use statistical significance tests to interpret simulation model results”. Oikos 123.4, pp. 385–388. DOI: 10.1111/j.1600-0706.2013.01073.x.

Wolfram Research, I. (2022). Mathematica. Champaign, IL: Wolfram Research, Inc. URL: https://www.wolfram.com/mathematica.

Zaehle, S. et al. (2014). “Evaluation of 11 terrestrial carbon–nitrogen cycle models against observations from two temperate Free-Air CO2 Enrichment studies”. New Phytologist 202.3, pp. 803– 822. DOI: 10.1111/nph.12697.

